# Impact of cell-type and context-dependent regulatory variants on human immune traits

**DOI:** 10.1101/2020.07.20.212753

**Authors:** Zepeng Mu, Wei Wei, Benjamin Fair, Jinlin Miao, Ping Zhu, Yang I Li

## Abstract

The effects of trait-associated variants are often studied in a single relevant cell-type or context. However, for many complex traits, multiple cell-types are involved. This applies particularly to immune-related traits, for which many immune cell-types and contexts play a role. Here, we studied the impact of immune gene regulatory variants on complex traits to better understand genetic risk mediated through immune cell-types. We identified 26,271 expression quantitative trait loci (QTLs) and 23,121 splicing QTLs in 18 immune cell-types, and analyzed their overlap with trait-associated loci from 72 genome-wide association studies (GWAS). We showed that effects on RNA expression and splicing in immune cells colocalize with an average of 40.4% and 27.7% GWAS loci for immune-related and non-immune traits, respectively. Notably, we found that a large number of loci (mean: 14%) colocalize with splicing QTLs but not expression QTLs. The 60% GWAS loci without colocalization harbor genes that have lower expression levels, are less tolerant to loss-of-function mutations, and more enhancer-rich than genes at colocalized loci. To further investigate the 60% GWAS loci not explained by our regulatory QTLs, we collected H3K27ac CUT&Tag data from rheumatoid arthritis (RA) and healthy controls. We found several unexplained GWAS hits lying within regions with higher H3K27ac activity in RA patients. We also observed that enrichment of RA GWAS heritability is greater in H3K27ac regions in immune cell-types from RA patients compared to healthy controls. Our study paves the way for future QTL studies to elucidate the mechanisms of as yet unexplained GWAS loci.

## Introduction

Genome-wide association studies (GWAS) have identified well over ten thousand genomic loci associated with human diseases and complex traits. However, while the number of trait-associated variants continues to grow, the causal genes and mechanisms at most GWAS loci remain difficult to determine. This difficulty is in part owing to the fact that ~90% of GWAS variants lie in noncoding regions [1].

Multiple studies have now shown that noncoding trait-associated variants are enriched for expression QTLs (eQTLs) and enriched in regulatory elements such as enhancers and promoters [2–4]. These findings suggest that noncoding variants likely affect traits by impacting gene regulation, which has motivated many studies to map regulatory QTLs – in particular eQTLs – in a diverse set of tissues and cell-types [5–12]. While eQTLs indeed overlap with many variants that have been associated with complex traits and diseases [2], several studies that assessed colocalization between GWAS and eQTL variants concluded that only a minority of GWAS loci can be explained by the eQTLs detected in available samples. For example, a 2017 study [13] reported that ~25% of variants associated with autoimmune disease colocalize with eQTLs in at least one of three immune cell-types they analyzed. In addition, a paper from the GTEx consortium suggests that ~20% of GWAS loci show colocalized effect with eQTLs in the tissue most relevant to the trait, and this number increases to an average of ~52% when all tissues were jointly considered [14]. Although the latter observation may suggest that genetic effects in very different cell-types together impact disease risk, a more recent study found that predicting causal genes based on eQTL colocalization with GWAS loci has a very low predictive value (just 17%) when eQTLs from all GTEx tissues were considered [15]. These data indicate that regulatory variants often impact the expression levels of different genes in different tissues, and imply that to minimize false positives, we should focus on determining colocalization in tissues or cell-types relevant to the trait under study. Moreover, another recent study estimated that only an average of 11% of trait narrow-sense heritability could be explained by *cis*-genetic effects on gene expression levels as measured in GTEx [16]. These data further suggest that to better understand the genetic basis of human traits, possible approaches are to (i) identify eQTLs in cell-types immediately relevant for each trait and (ii) to expand the study of genetic effects on gene regulatory mechanisms beyond that of steady-state mRNA expression levels alone.

The goal of this study was to better understand how regulatory variants in immune cell-types affect common disease risk. To this end, we performed a uniform eQTL and sQTL mapping in four datasets, including (i) a dataset with a large number of different immune cell-types but a small sample size (DICE [7]), (ii) a dataset with a single tissue-type but with a large sample size (DGN [9]), and (iii) two intermediate datasets (BLUEPRINT [6], and GEUVADIS [8]). In particular, the DICE consortium collected population RNA-seq data for 13 unstimulated immune cell-types including various naïve and effector/memory T cell subtypes, classical and non-classical monocytes, B cells, and NK cells. DICE also performed *in vitro* activation of CD4^+^ and CD8^+^ T cells using CD3/CD28 antibodies. Although, the sample size in the DICE dataset is the smallest among the four datasets (~90), the large number of sorted cell-types makes the DICE dataset ideal to identify cell-type-specific genetic effects. The BLUEPRINT consortium collected data for three cell-types in ~197 individuals, classical monocytes, naïve CD4 T cells and neutrophils, the last of which was not collected by the DICE consortium. The DGN dataset include 922 whole blood samples and, finally, the GEUVADIS dataset consists of RNA-seq data from 462 lymphoblastoid cell lines (LCL), which are derived from B-cells. We reasoned that analyzing these datasets in a uniform fashion (**Figure 1a**) would allow us to capture both strong eQTLs with cell-type-specificity (using DICE) and weak-effect eQTLs that are less likely to be cell-type-specific (e.g. using DGN).

**Figure 1.**
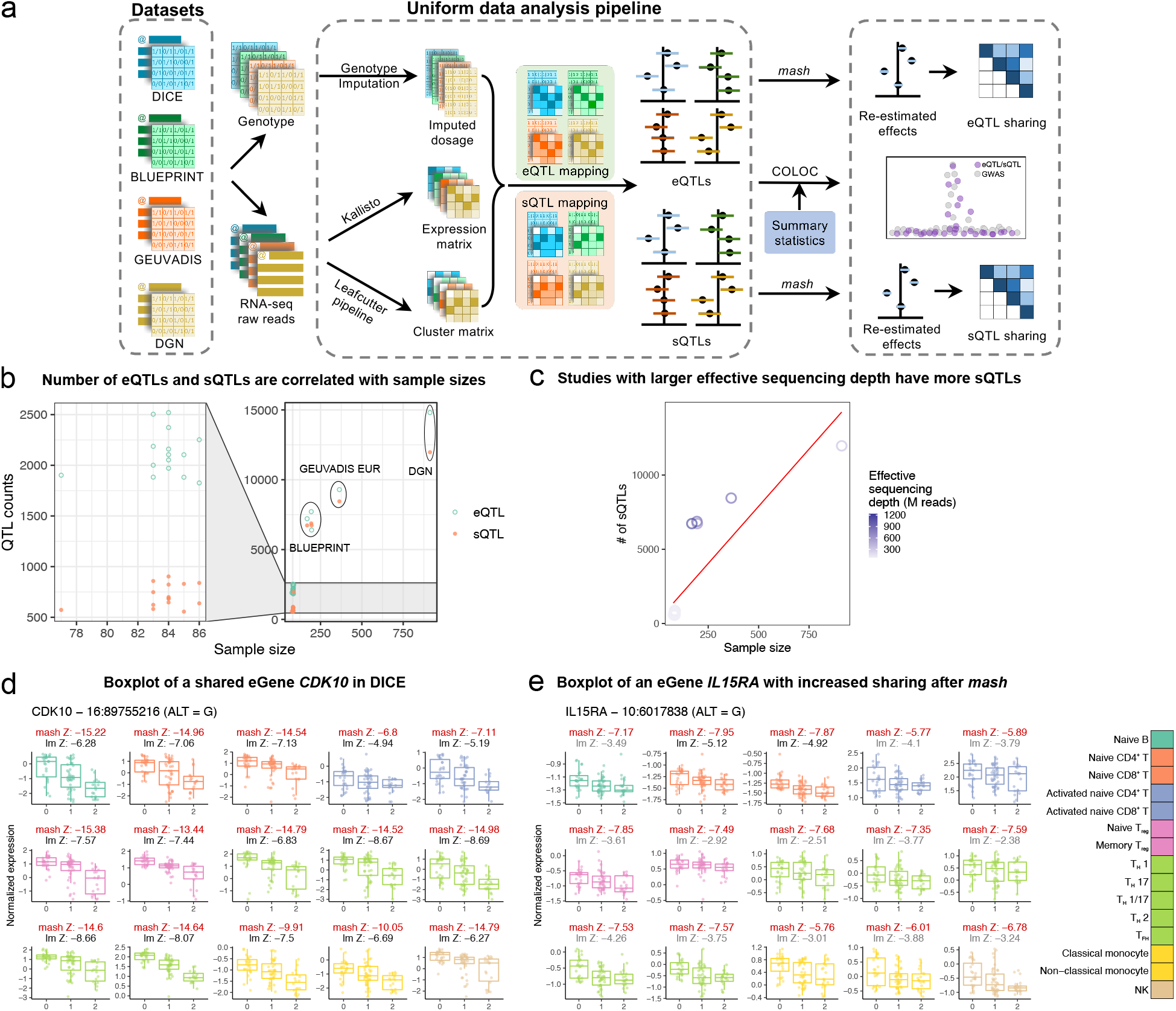
Summary of analysis workflow. **(a)** A uniform computational pipeline to analyze data from four large immune RNA-seq datasets (DICE, BLUEPRINT, GEUVADIS, and DGN). The same pipeline for genotype imputation, expression and splicing quantification and QTL-mapping were applied to the four datasets. Sharing of QTLs among cell-types were quantified using *mash* [18]. Colocalization analyses were performed for 72 GWAS of immune-related and non-immune traits. **(b)** Total number of genes and intron clusters with a significant QTL identified in DICE (left) and the other three studies (right) as a function of sample sizes. QTLs are considered significant when Storey’s q-value is below 0.05. **(c)** Studies with larger effective sequencing depth (BLUEPRINT and GEUVADIS EUR) have more sQTLs comapred to other studies. Effective sequencing depth = library size × read-length. Red line represents the fitted line in a simple linear model. **(d)** An eQTL at the gene *CDK10* that is shared by all 15 cell-types in DICE despite large differences in baseline expression levels across cell-types. **(e)** An eQTL at the *IL15RA* gene that is shared across immune cell-types but show cell-type-specificity according to linear regression. lm Z: Z-scores of linear model from `FastQTL`, mash Z: Z-scores estimated by *mash* (red). The lm Z-scores were colored in grey when the Z-score did not pass statistical significance after `FastQTL` permutation and in black when they were determined to be significant.

Using these uniformly processed QTLs, we found that, on average, eQTLs and sQTLs together colocalize with 40.4% GWAS loci for autoimmune diseases and blood-related phenotypes, doubling for many GWAS the number of colocalizing loci from previous studies [13]. To further study the biology underlying GWAS loci, we collected H3K27ac profiles of 5 immune cell-types (CD4^+^ T cells, CD8^+^ T cells, regulatory T cells, monocytes, and B cells) from rheumatoid arthritis (RA) patients and healthy controls using CUT&Tag [17]. These additional data helped us to better understand the cellular context in which GWAS loci, including those without colocalization, contribute to disease risk. Taken together, our work reports a comprehensive analysis of the regulatory effects of genetic variants on immune cell-types and their overlap with GWAS loci and with gene regulatory regions as measured in a disease cellular context. Our maps of eQTLs, sQTLs, and gene regulatory regions in diverse immune cell-types are available online, which we foresee will aid research on the genetic basis of diseases and on gene regulation in immune cells.

## Results

### Harmonized map of eQTLs and sQTLs in 18 immune cell-types

We built a uniform data processing pipeline to harmonize four large population-scale RNA-seq datasets (**Figure 1a**). Our pipeline, which is described in detail in the Methods section, includes quantifying RNA expression and splicing levels, imputing genotype data to the same common reference panel, and calling QTLs in all datasets using the same methods. We also designed an approach to harmonize quantification for splicing junction usage across cell-types and datasets by first merging `LeafCutter` intron clusters across all samples and then re-calculating intron usage for each sample (Methods). Thus, our approach produced a harmonized set of introns that can be readily interrogated.

To map expression QTLs (eQTLs) and splicing QTLs (sQTLs), we used `FastQTL` [19]. As covariates for the linear regression, we used three genotype PCs and a number of phenotypic PCs chosen to maximize the number of significant QTLs (Storey’s q-value < 0.05) (**Supplementary Table**). We used `LeafCutter` [20] to collapse introns that share a splice site into intron clusters, as genetic effect on multiple introns within a cluster tend to reflect the impact of a single causal genetic variant [20]. In total, we discovered 26,271 genes and 23,121 intron clusters that have a significant QTL in at least one the four datasets. As expected, both the numbers of eQTLs and sQTLs were correlated with sample size (**Figure 1b**). In addition to the sample size, we found that the number of sQTLs identified was also correlated with effective sequencing depths (**Figure 1c**). For example, while the number of sQTLs is roughly linearly related to sample size, datasets with higher effective sequencing depths consistently yielded more sQTLs than predicted by a simple linear model. This is most obvious for BLUEPRINT, which used 100bp single-end or paired-end sequencing when compared to DICE or DGN (both 50bp single-end).

We show a shared eQTL for the *CDK10* gene (**Figure 1d**) and an eQTL for the *IL15RA* gene (**Figure 1e**) as examples. All gene expression and splicing quantifications are available as Supplementary Files (**Supplementary File 1** and **Supplementary File 2**), including all identified expression and splicing QTLs (**Supplementary File 3**).

### Global patterns of eQTL and sQTL sharing across immune cell-types

To characterize the cell-type-specificity of genetic effects in immune cells, we sought to discern genetic variants that impact gene regulation broadly across many or all immune cell-types from those that impact a few or only one cell-type. Previous studies have also quantified the sharing and specificity of regulatory QTLs [6, 7]. However, because the sample sizes of most datasets are small, we speculated that estimates of QTL effect sizes are noisy, which would generally cause studies to underestimate the levels of QTL sharing.

We reasoned that our harmonized dataset would allow us to better infer sharing patterns. In particular, we improved our estimates of eQTL and sQTL effect sizes at each locus by using a multivariate adaptive shrinkage method *(mash)* [18]. The *mash* method improves estimates of QTL effect sizes from those that are obtained from applying linear regression in each cell-type separately, because *mash* leverages the correlation structure of QTL effect sizes across all cell-types to re-estimate QTL effect sizes at each locus. To use *mash*, we first identified 36,950 unique SNP-gene associations and 116,881 unique SNP-intron associations (q-value below 5%) in the 15 DICE cell-types separately. We built a *mash* model from all protein-coding genes for eQTLs and all introns for sQTLs, and applied it to calculate the posterior mean effect sizes, henceforth referred to as *mash* effect sizes, for the 36,950 SNP-gene and 116,881 SNP-intron pairs in each of the 15 cell-types. This procedure increased the effective sample sizes approximately 8- to 12-fold for gene expression (28- to 40-fold for RNA splicing) as estimated by *mash* (**Supplementary Figure 2**, see Methods), and thus greatly enhanced estimates of QTL effect sizes in the 15 immune cell-types (for two examples see Z-scores in **Figure 1d, e**).

We first asked about the proportion of QTLs that are shared across immune cell-types based on the estimated *mash* effect sizes. We found that a large fraction (33.7%, n = 2,897 of 8,597) of genes with an eQTL (eGenes) are shared according to *mash* (LFSR < 0.05) across all six distinct major cell-types collected in the DICE dataset (B cell, naïve CD4^+^ and CD8^+^ T cell, NK cell, classical monocytes, non-classical monocytes) (**Figure 2a**). Our estimates of sharing are therefore much higher than the 5.2% (463 out of 8,863) of sharing estimated in the original DICE study [7]. In fact, the original DICE study estimated that nearly half of all eGenes are specific to a single immune cell-type, while our new estimate suggests that only 20.4% are likely cell-type-specific (**Figure 2a**).

**Figure 2.**
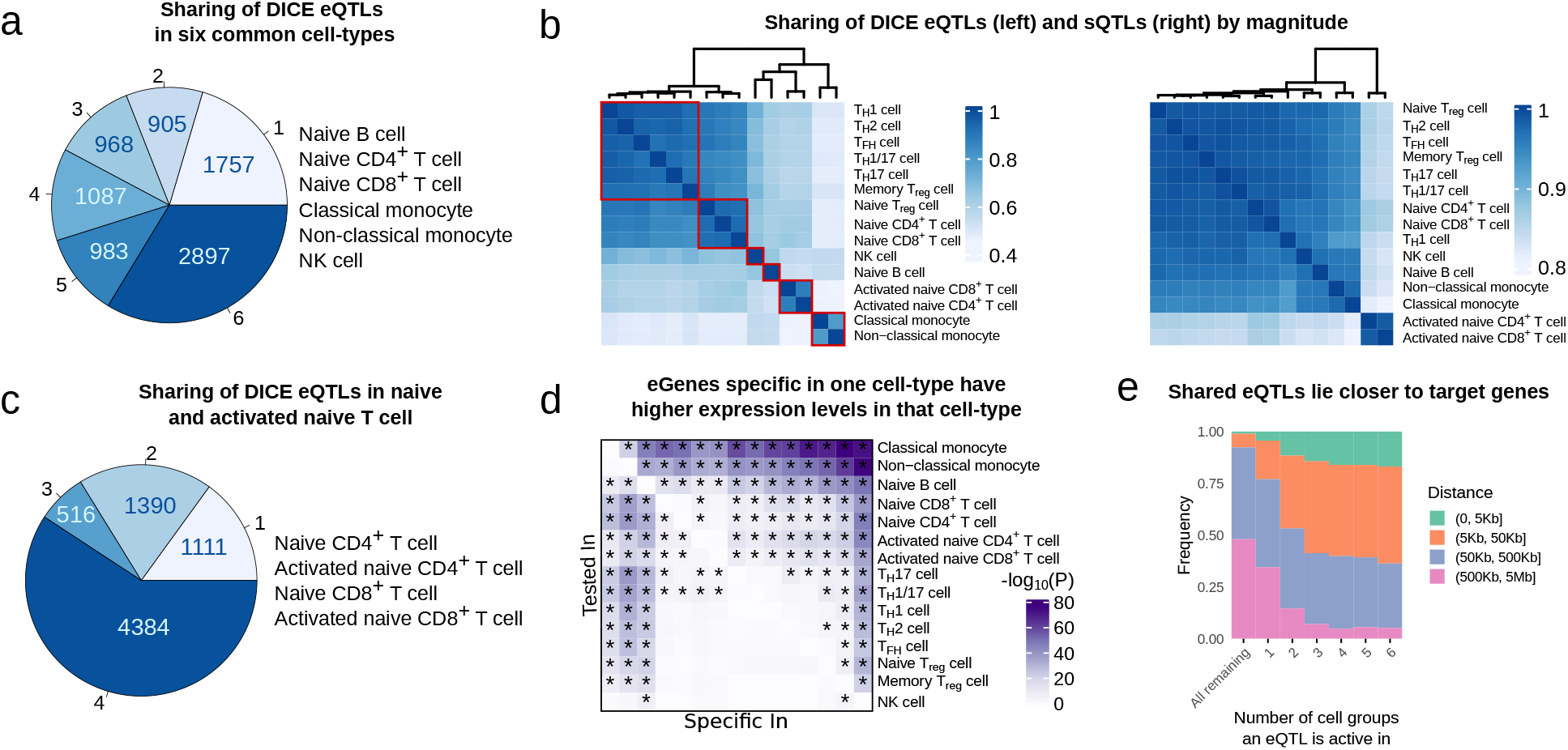
Sharing of eQTLs and sQTLs using *mash*. **(a)** Estimated number of cell-types in which eQTLs are inferred to be active out of six DICE cell-type groups according to *mash*. These estimates of sharing are much higher than that from the original study [7]. **(b)** Sharing of eQTLs (left) and sQTLs (right) by magnitude. Red square: cells were grouped into six clusters based on eQTL sharing, which resulted in the following groups: (i) Naïve T cells, (ii) Memory and Effector T cells, (iii) Monocytes, (iv) activated T cells, (v) B cells, and (vi) NK cells. **(c)** Estimated number of cell-types in which eQTLs are inferred to be active among naïve and activated naïve T-cells from DICE. **(d)** Heatmap showing −log_10_ p-values of a differential gene expression analysis that compared the expression level of cell-type-specific eGene in the discovery cell-type to the expression levels of the eGene in the other 14 cell-types. **(e)** Distance between eQTLs and their target genes stratified by the number of cell groups in which the eQTL is active. To obtain the six cell groups, we grouped the 15 cell-types based on similarity in their *mash* effect sizes as described in **(b)**.

Using *mash* effect sizes, it is also possible to quantify the amount of QTL sharing in terms of magnitude of effects. We found that over 40% of eQTLs have similar *mash* effect sizes (within 2-fold, 34% within 1.5-fold) across pairs of cell-types, and this fraction increases to over 90% when considering closely related cell-types, such as classical and non-classical monocytes or T_H_1 and T_H_17 cells (**Figure 2b**, left). In addition, we found that the vast majority (> 80% within 2-fold, 67% within 1.5-fold) of sQTLs have similar *mash* effect sizes across all immune cell-types, with activated CD4^+^ and CD8^+^ T cells forming an outlier group in a hierarchical clustering based on estimates of sharing (**Figure 2b**, right). These results are consistent with previous work [5] and suggest that the impact of genetic variation on RNA splicing is generally shared across two celltypes when the involved mRNA transcripts are expressed in both cell-types, which is largely the case for any pair of immune cell-types.

In general, the proportion of shared eQTLs across cell-types captured the lineage relationships among the 15 immune cell-types. Specifically, classical and non-classical monocytes clustered together, while B cells and NK cells each formed distinct clusters. Furthermore, despite a high level of QTL sharing (> 80%) among naïve T cells, we found that naïve CD4^+^, CD8^+^ and regulatory T cells formed one cluster, while memory and effector T cells formed another larger cluster. We also observed a higher level of QTL sharing between activated CD4^+^ and CD8^+^ T cells compared to that between stimulated and naïve T cells. This observation suggests that activated CD4^+^ and CD8^+^ T cells share similar gene expression programs upon activation, and that differences in genetic effects on gene regulation exist between activated and non-activated cells. Nevertheless, we found that 66.2% of eGenes (n = 4,900) were shared according to *mash* (LFSR < 0.05) between naïve and activated T cells, suggesting that the overall impact of genetic effects on gene regulation is in most cases the same across activated and naïve T cells (**Figure 2c**).

While a large proportion of eQTLs and sQTLs appeared to be shared across multiple immune cell-types, we found that a substantial number of eQTLs (2810, 27.8%) appeared cell-type-specific. We asked whether QTLs that appeared cell-type-specific showed specific features compared to QTLs that were shared across immune cell-types. We first asked whether genes with eQTLs that were specific to a cell-type were also more highly expressed in that cell-type compared to the other cell-types. To test this, we asked whether genes with an eQTL in a cell-type *A* but not in another cell-type *B,* were significantly more highly expressed in cell-type *A* than cell-type *B.* Indeed, we found that this was the case for most cell-type-specific eQTLs (66.7%, Bonferoni adjusted P-value < 0.05, one-sided, paired Wilcoxon rank-sum test), suggesting that variation in gene expression level likely impacts whether a genetic variant has a regulatory effect and/or our ability to detect this effect. This observation was most obvious for classical monocytes, non-classical monocytes and naïve B cells, and is driven by differences in their gene expression levels compared to T cells (**Figure 2d**). In addition to differences in gene expression levels, we found that eQTLs that were cell-type-specific were located further away from the gene transcription start site in comparison to eQTLs that were shared across immune cell-types (**Figure 2e**). Moreover, cell-type-specific eQTLs were more highly enriched in enhancers compared to eQTLs that were shared (**Supplementary Figure 3**). These observations are consistent with the notion that cell-type-specific eQTLs tend to impact enhancer activity, while shared eQTLs more often impact promoters [21].

Taken together, our analyses revealed that QTL effects are shared for a large number genes. Nevertheless, we were able to detect a non-negligible number of cell-type or cell-group specific QTLs. Importantly, these findings and classification show replication across datasets (Supplementary Note 1). Thus, we expect our QTL data to be highly replicable in existing or future immune QTL datasets.

### Colocalization of immune regulatory QTLs with common disease GWAS

Our harmonized eQTL and sQTL data gave us the unprecedented ability to identify genetic variants that impact traits through regulatory effects on immune cell-types. We performed colocalization analyses that aimed to determine whether the genetic variants at GWAS loci that are causal for a trait are likely to be the same variants as the causal regulatory QTLs. We compiled a set of 72 well-powered GWAS, including 14 for autoimmune diseases (11 unique disease types), 36 blood traits, and 22 other traits (**Supplementary Table**), and used `COLOC` to evaluate colocalization (PP4 ≥ 0.75) [22] with DICE, BLUEPRINT, and DGN QTLs separately (**Supplementary File 4**; average N = 206,090). We report the main colocalization results of our analyses using BLUEPRINT QTLs (3 immune cell-types) below, and use the DICE (15 cell-types) regulatory QTLs to interpret the cell-type-specificity of colocalized genes. We reasoned that choosing BLUEPRINT over DICE as the main dataset for this analysis will increase our power for QTL mapping owing to its larger sample size and will also allow us to identify more sQTLs owing to higher RNA-seq coverage and longer read lengths (**Figure 1c**).

When we ascertained colocalization between GWAS loci for the 72 traits and QTLs from BLUEPRINT, we observed that colocalization rates between immune regulatory QTLs and GWAS hits were higher for autoimmune and blood-related traits compared to other non-immune traits (mean 40.4% versus 27.7%) (**Figure 3a**). This observation supports the expectation that a large fraction of colocalized regulatory QTLs indeed affect immune traits by impacting gene regulation in immune cell-types.

**Figure 3.**
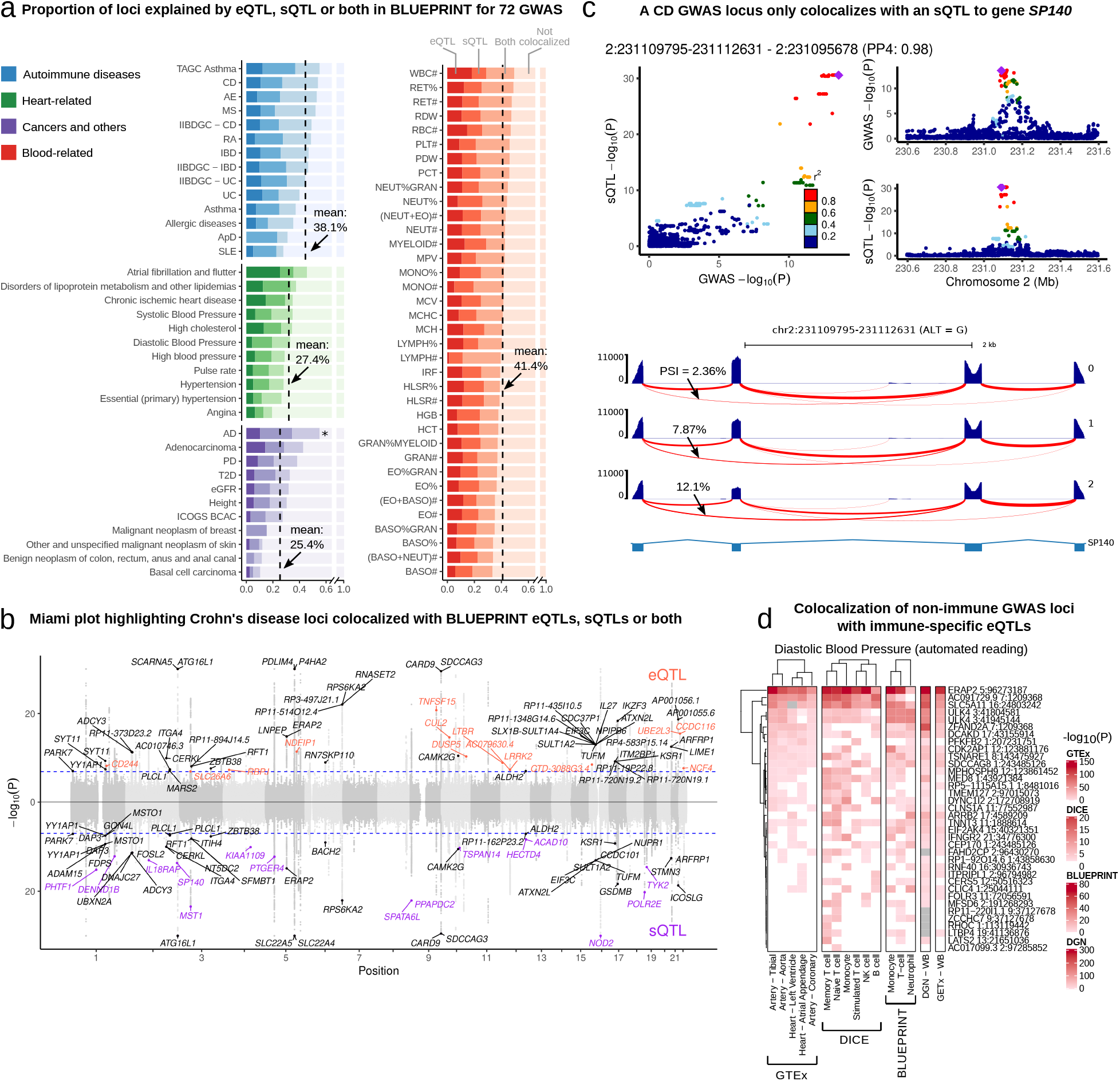
Colocalization analysis explained up to 47% of GWAS variants and revealed potential causal SNPs to non-immune traits. **(a)** Proportions of GWAS loci colocalized with eQTLs, sQTLs, or both. Dashed line: mean colocalization rate. *: Alzheimer’s Disease (AD) GWAS was not included in the mean calculation owing to the well-documented involvement of microglia in AD. **(b)** Colocalization of Crohn’s Disease (CD) GWAS with eQTLs (orange), sQTLs (purple), or both (black). GWAS SNPs with −log_10_(P) larger than 30 were set to 30 to facilitate visualization. **(c)** LocusCompare plot (top) and Sashimi plot (bottom) showing colocalization between a sQTL of an intron in gene *SP140* in T cell and a GWAS locus for CD. Arrows in the Sashimi plot point to the intron affected by the sQTL, labeled with PSI quantification from `LeafCutter` [20]. **(d)** Heatmap of eQTL association strengths in GTEx tissues and immune celltypes for DICE eQTLs that colocalize with diastolic blood pressure loci. Several GWAS loci colocalize with eQTLs active in immune cell-types but not in heart-related tissues.

We next focused on autoimmune diseases and blood-related traits. Our regulatory QTLs colocalized with a mean of 38.1% (range: 24–47.4%, n = 14) and 41.4% (range: 33.8–50%, n = 36) of autoimmune disease and blood traits GWAS loci (PP4 > 0.75), respectively (**Figure 3a**). The mean rates of colocalization ranged from 27.4% to 50.2% depending on the choice of posterior probability cutoff for determining colocalization status (PP4, ranging from 0.5 to 0.9, **Supplementary Figure 5**). We chose to use an intermediate cutoff of 0.75 to be consistent with previous studies [13]. Expression QTLs colocalized with 6.7-39.3% of GWAS loci with an average of 26.6%, similar to estimates from previous studies [13]. Notably, we found that splicing QTLs colocalized with an additional 7.6–21% GWAS loci (average: 13.8%) that did not colocalize with an eQTL, and explain much of the increase in colocalization rates from this study compared to that of previous studies. Interestingly, we observed that the rates of colocalization between GWAS loci and both an eQTL and an sQTL can vary substantially across traits, ranging from 4.5% for systemic lupus erythematosus (SLE) to 28.6% for basophil percentages of granulocyte (BASO%GRAN) (average: 17.2%). Most notably, nearly all colocalized loci associated with SLE (10 out of 16) colocalized only with sQTLs (**Figure 3a**). This result raises the possibility of distinct regulatory architectures for different diseases. We obtained similar rates of colocalization with DICE and DGN, for which 30.7% and 38% immune GWAS loci colocalized with DICE and DGN regulatory QTLs, respectively (**Supplementary Figure 7**).

To help the interpretation of these results, we show the colocalizations between immune regulatory QTLs and GWAS loci for Crohn’s disease (CD) as an example [23] (**Figure 3b**) (12,194 cases and 28,072 controls). We included 108 GWAS loci in our colocalization analysis that pass a p-value threshold of 10^−7^ (see Methods). Ten and fifteen loci colocalized with only eQTLs or sQTLs, respectively, while an additional 25 loci colocalized with both eQTLs and sQTLs. In total, 46% of loci colocalized with an eQTL, an sQTL, or both. Of note, several identified colocalized genes have been extensively studied in terms of CD etiology, including *NOD2* [24] and *ITGA4,* of which the latter is the target for the CD monoclonal antibody drug natalizumab [25].

The high rates of colocalization (average: 13.8%) between GWAS loci and sQTLs highlight the importance of considering the impact of risk variants on RNA splicing. For example, we identified an sQTL associated with the skipping of the seventh exon in gene *SP140* in T cells that colocalized with a risk locus in both CD GWAS we analyzed (**Figure 3c**) [23, 26]. *SP140* encodes nuclear body protein SP140 [27], which preferentially binds to gene promoters with H3K27me3 modification [28] and regulates multiple immune-related genes [29]. Notably, the exclusion of the same exon in *SP140* transcript isoforms has also been associated with risk alleles for other diseases including multiple sclerosis [30].

Immune regulatory QTLs colocalized at a lower rates in GWAS of traits that are not autoimmune or blood-related (27.7%). Among the 22 non-immune traits we analyzed, Alzheimer’s disease (AD) is an outlier, for which 55% of GWAS loci colocalized with a BLUEPRINT QTL. The high rate of colocalization can be explained by the known role of microglia in AD etiology [31]. To make sense of the colocalizations found in GWAS of other non-immune traits, we reasoned that colocalized loci may either reflect a causal effect of the risk variant on disease through immune cell-types, or a causal effect of the risk variant on disease through non-immune cell-type but that is also manifested in an immune cell-type. However, if a regulatory QTL effect for the GWAS locus is only detected in an immune cell-type, then it is more likely that the GWAS variant impacts the trait through immune cell-types.

Because immune cells are well known to play a role in many human traits including non-immune traits (see Supplementary Note 2), we sought to identify trait-associated variants that likely act through immune cell-types. To this end, we obtained association statistics between all colocalized eQTL SNPs (in DICE) and the gene expression level in GTEx tissues, and asked about the proportion of colocalized eQTLs that do not show any effects on gene expression levels in tissues that are most relevant to each trait. We chose five GTEx heart tissues as the relevant tissues for our 11 heart-related GWAS. For breast cancer, we ascertained the effect of colocalized eQTL SNPs in breast and adipose tissues. For Parkinson’s disease (PD), we used the 13 GTEx brain tissues. Overall, we found that 65 of 267 (24.3%) loci that colocalized in the 14 selected GWAS are specific in DICE and BLUEPRINT immune cell data (**Supplementary File 5**, Supplementary Note 3). For example, we found that 8 of 36 eQTLs that colocalized with diastolic blood pressure in DICE are not significant in any of the five heart related tissues (P-value > 0.05 / 5) (**Figure 3d**). Of note, two genes with colocalized eQTLs, *FOLR3* and *LTBP4*, were significantly associated with gene expression levels in GTEx whole blood, suggesting that they indeed likely function through the immune cells. Interestingly, this possibility is further supported by studies that have shown that *FOLR3* is down-regulated in peripheral blood of patients with hypertension [32, 33].

### GWAS-eQTL colocalizations across immune cells are highly shared when accounting for statistical power

Several studies have proposed that a large fraction of autoimmune disease risk loci affect gene expression levels in a cell-type-specific manner [34, 35]. We sought to use our dataset to evaluate this hypothesis by analyzing the cell-type-specificity of the eQTLs that colocalize with autoimmune GWAS loci. For this, we focused on the 197 genes with a DICE eQTL (eGenes) that colocalized with at least one of the 14 autoimmune disease GWAS in our study. We then evaluated the celltype-specificity using the *mash* QTL effect sizes estimated for the 15 immune cell-types from the DICE consortium.

The general pattern of sharing that we observed for colocalized risk loci is that the corresponding eGenes are mostly shared across multiple cell-types. The sharing was also apparent across the 6 major groups of immune cells that represent naïve T cells, memory and effector T cells, monocytes, activated T cells, B cells, and natural killer (NK) cells (**Figure 4a**). Remarkably, 65 of 197 (33.0%) tested genes colocalized in all 6 major immune cell groups. The immune cell groups in which the most colocalized genes were found are memory and naïve T cells, in which 160 and 151 of 197 eGenes colocalized with GWAS loci, respectively. Only 8, 8, and 4 eGenes showed an effect that appear to be specific to B cells, monocytes, and NK cells, respectively, while 12 eGenes show an effect only in T cells. These observations suggest that for the vast majority of autoimmune risk loci, the effect of risk variants on gene expression level is not restricted to a single immune cell-type or cell group.

**Figure 4.**
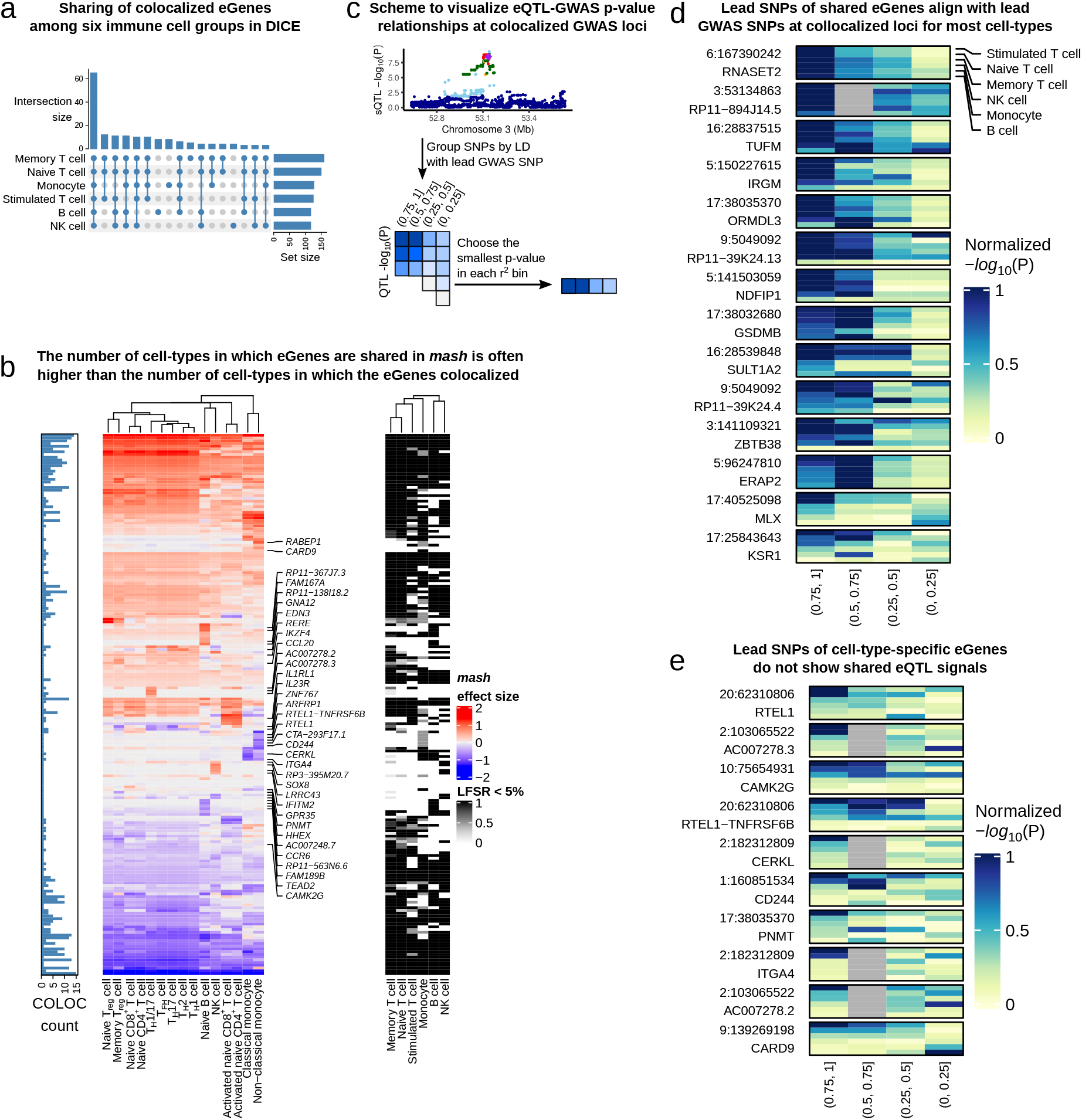
*mash* analysis indicates high sharing of QTLs among immune cell-types. **(a)** Upset plot showing the cell-group-specificity or sharing of eGenes colocalized with immune-related GWAS loci. The majority of colocalized eGenes are shared across the 6 cell groups. **(b)** Heatmaps showing *mash* effect sizes of colocalized eGenes (left) and LFSR (< 0.05, right). Barplot on the left shows the number of cell-types in which the eGenes was determined to colocalize with a GWAS variant using COLOC. While the mash effect sizes are estimated to be shared across most immune cell-types for most GWAS loci, the colocalization status as determined using `COLOC` (PP4 > 0.75) often imply cell-type-specificity. **(c)** Schematic representation of our approach to visualize the QTL association P-value distribution of colocalized eGenes across SNPs with different amount of LD with the lead GWAS SNP. If a QTL in a cell-type colocalizes with a GWAS loci, then in general the significance of the QTL association should decrease for SNPs with decreasing amount of LD with the lead GWAS SNP. **(d)** eQTL p-values in different LD bins (as described in **(c)**) at GWAS loci with colocalized eQTLs across all 6 cell-groups. Colocalized eGenes that were inferred to be shared all have lower eQTL p-values at SNPs in high LD with the lead GWAS SNPs. **(e)** By contrast, colocalized eGenes that were inferred to be cell-type specific show different patterns of eQTL p-value distribution in the LD bins.

We next set to understand the discrepancy between our finding that most GWAS loci impact multiple cell-types and that of previous work, which suggests more cell-type-specificity [34, 35]. We first analyzed colocalization status of autoimmune GWAS loci in each cell-type separately, which corresponds to the general approach used by previous studies [5, 7, 13]. We found that, using this approach, the number of cell-types with positive colocalization status is generally smaller – sometimes much smaller – than the number of cell-types in which the eQTL effects are shared according to our analysis (**Figure 4b**). In other words, the number of cell-types in which a GWAS locus colocalizes with an eQTL is generally smaller than the number of cell-types in which that same eQTL is inferred to be active. We speculated that this discrepancy results from the variation in the posterior probabilities of colocalization computed by `COLOC`, owing to inherent noise in estimating the effect sizes and statistical significance of eQTLs. In support of this, we found a gene *RNASET2,* whose eQTLs colocalized with a CD risk locus in 7 out of the 13 cell types analyzed (PP4 ranges between 0.79 and 0.99), but whose eQTLs were inferred to be active across all 13 cell-types (**Supplementary Figure 11**). We found that in the colocalized cell-types, the lead eQTL SNP was also the lead GWAS SNP. In the 7 other cell-types, the lead eQTL SNPs did not correspond exactly to the lead GWAS SNP, but were in strong LD (r^2^ > 0.6). As a result of this variation, the posterior probabilities of colocalization (PP4 values) in these 6 cell-types ranged from 0.58 to 0.69, which did not pass our cutoff of 0.75. Taken together, these observations suggest that *RNASET2* eQTLs colocalize with the Crohn’s disease GWAS locus in all 13 immune cell-types.

We next asked whether this observation was reflective of a general trend across GWAS loci. We reasoned that, under the simplifying assumption that there is only one causal eQTL at each GWAS locus, colocalized loci should show a general pattern where SNPs in high LD with the lead GWAS SNP will show strong associations with expression levels of the colocalized gene, but the eQTL associations will weaken for SNPs in lower LD. Thus, eQTLs that colocalize with a GWAS locus in all cell-types should show decreasing eQTL association strength for SNPs in decreasing amount levels of LD for most or all cell-types. By contrast, eQTLs that only colocalize with a GWAS locus in a single cell type, should show these patterns only in a single or a small number of cell-types.

To visualize these patterns across many GWAS loci and cell-types, we first found the lead GWAS SNPs at every colocalized loci and divided all SNPs within 1Mb into four bins according to their linkage disequilibrium (LD) with the lead SNP (namely, r^2^ within ranges of (0, 0.25], (0.25, 0.5], (0.5, 0.75], and (0.75, 1]). Next, for each r^2^-bin, we identified the SNP with the smallest eQTL p-value for the colocalized eGene in each of the 6 DICE cell-groups (**Figure 4c**). We then plotted the p-values for all the colocalized locus-gene pairs where the *mash* SNPs and the lead GWAS SNPs are in close LD (r^2^ > 0.8, **Figure 4d**). We observed that the top eQTLs are often in high LD with the lead GWAS SNP for multiple cell-groups (rows) when the eQTLs were determined to have shared effects. By contrast, for the eQTLs we inferred to have a cell-type-specific effect, the patterns are strikingly different as the most significant eQTLs are more likely to be in lower r^2^-bins in most cell-types (**Figure 4e**). These findings support our high estimation of shared regulatory effects of GWAS variants across multiple cell-types. These observations also suggests that `COLOC` is susceptible to noise in QTL mapping, especially when the sample size in QTL mapping is small. More importantly, our data indicates that previous work likely over-estimated the number of GWAS loci with cell-type-specific colocalization with eQTLs.

Overall, we find evidence that previous work overestimated cell-type-specificity of GWAS effects on gene expression levels. Indeed, using our data and analysis methods, we found cell-type-specific colocalization in a single or two major immune cell-groups for only 35 of 197 loci (17.8%). By contrast, we found that up to 103 (52.3%) GWAS loci are eQTLs in five or more cell-groups.

### Limited regulatory effects specific to stimulated cells at GWAS loci

Our analysis so far indicates that about 40% of autoimmune GWAS loci have a detectable effect on gene regulation in at least one of the 18 immune cell-types analyzed. We next wondered about the mechanism by which the remaining 60% of GWAS loci function. There are several possible explanations for why such a large fraction of GWAS loci do not colocalize with a regulatory QTL identified in our study. One simple explanation is that many of these GWAS loci do not impact disease risk by affecting the expression or splicing of mRNA. Instead, they may affect protein coding sequence or other as yet poorly studied molecular mechanisms, such as alternative polyadenylation [37].

To identify putative mechanisms by which trait-associated variants at uncolocalized GWAS loci function, we asked whether genes in GWAS loci without colocalization were different in terms of expression levels, enhancer density, and sequence constraint compared to those in GWAS loci with colocalization (Methods). Our analysis revealed that genes in loci without colocalization are expressed at a significantly lower levels than compared to genes at loci with colocalization (**Figure 5a**). In addition, we found a higher enhancer density as measured by EDS [38] (**Figure 5b**), and a lower tolerance to loss of function mutations as measured by LOEUF [36] (**Figure 5c**) for genes in uncolocalized GWAS loci.

**Figure 5.**
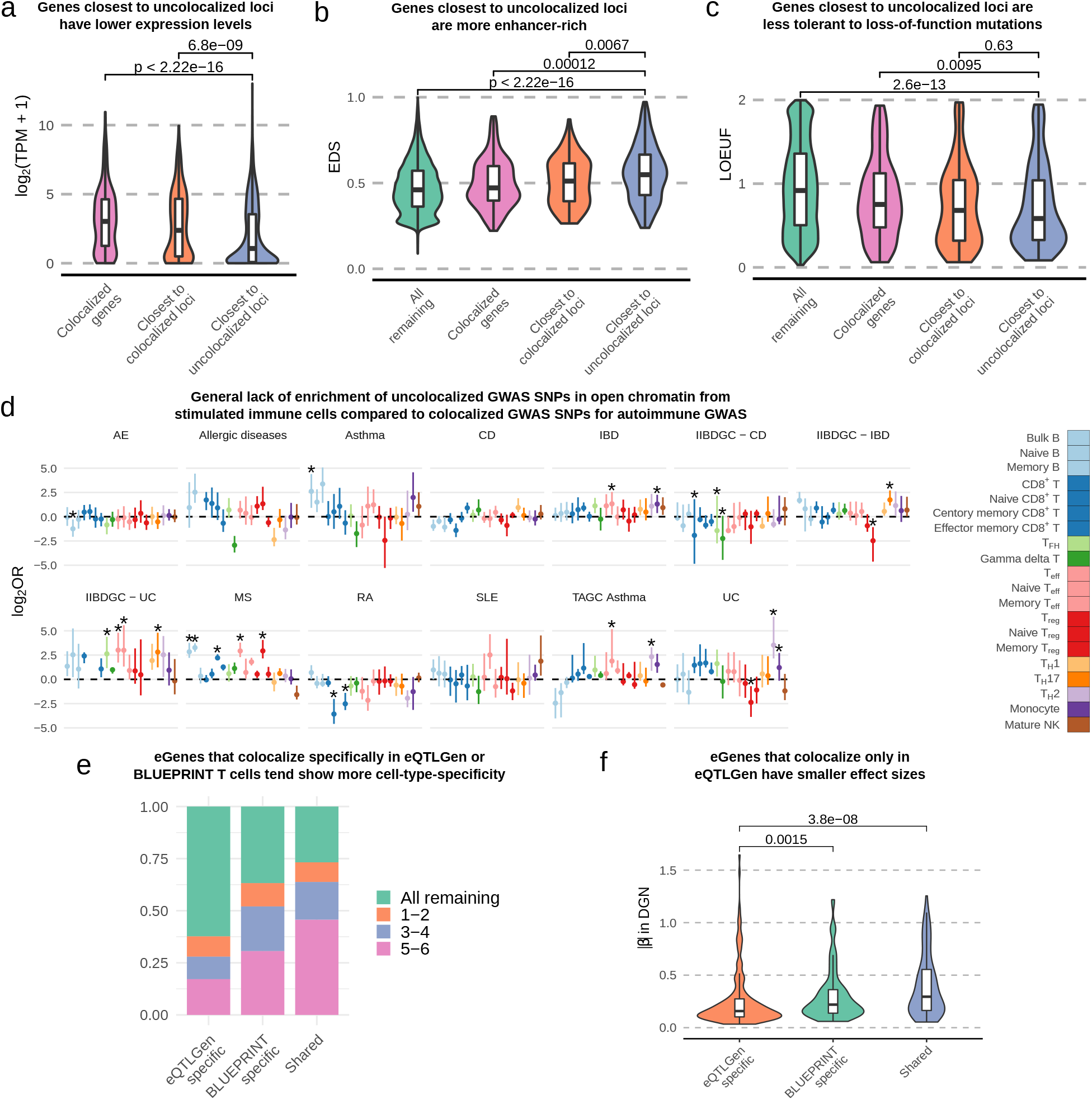
Characterizations of uncolocalized GWAS loci. **(a)** Genes closest to uncolocalized loci are expressed at lower levels compared to colocalized eGenes. **(b)** Genes closest to uncolocalized loci have higher EDS, indicating their expression is more constrained. **(c)** Genes closest to uncolocalized loci have lower LOEUF [36], suggesting that they are less tolerant to rare mutations. **(d)** Forest plot showing the log_2_ odds ratios of the enrichment of uncolocalized GWAS SNPs in open chromatin of stimulated immune cells compared to colocalized GWAS SNPs (Fisher’s Exact Test). Few stimulated immune cell-types are enriched for uncolocalized autoimmune GWAS loci. Error bars show 95% confidence intervals from bootstrap (Methods); *: FDR < 0.05. **(e)** eGenes that colocalized only in eQTLGen data tend to be restricted in fewer cell-types (in DICE data) compared to eGenes that colocalized only in BLUEPRINT data or eGenes that colocalized in both BLUEPRINT and eQTLGen. **(f)** eGenes that colocalize only in eQTLGen data have smaller effect sizes compared to eGenes that colocalize only in BLUEPRINT T cells or eGenes that are shared between eQTLGen and BLUEPRINT. To avoid the Winner’s curse effect, the effect sizes were ascertained using the DGN dataset.

Several studies have proposed that many autoimmune disease GWAS loci impact gene regulation in stimulated but not naïve immune cells [39, 40]. Thus, it is possible that a large fraction of uncolocalized GWAS hits impact gene regulation in stimulated but not unstimulated cells. However, we found in an earlier analysis that although some exception exist (**Supplementary Figure 12**), regulatory effects in stimulated CD4^+^ and CD8^+^ T cells were largely the same in unstimulated T cells. As a less direct but complementary analysis, we therefore asked whether uncolocalized GWAS loci were more likely to overlap with open chromatin regions in stimulated immune cells compared to colocalized ones using ATAC-seq data from 20 naïve and stimulated immune cells [39]. Again, however, we found very little support for the hypothesis that a large fraction of uncolocalized GWAS loci impact gene regulation in immune cells that were stimulated. Specifically, we observed very subtle differences in the enrichment of uncolocalized GWAS SNPs in open chromatin regions of stimulated immune cell-types compared to that of colocalized GWAS SNPs (**Figure 5d**, Methods). When accounting for multiple testing, only 17 out of 254 tests are significant at a FDR of 5%, and the enrichment for these were modest. Thus, these analyses suggest that there are fundamental differences in the mechanisms and genes that underlie colocalized and uncolocalized autoimmune GWAS loci, but the difference cannot be simply explained by regulatory effects that are restricted to stimulated immune cells.

Another explanation for the large number of uncolocalized GWAS loci is that the regulatory effects of many GWAS loci are outside current range of detection owing to small sample sizes. As a simple way to test this, we performed an eQTL analysis for only the lead GWAS SNPs at uncolocalized CD GWAS loci in BLUEPRINT T cells. The smaller number of tests compared to a genome-wide analysis improved our ability to detect eQTLs with smaller effect sizes (mean absolute effect size 0.34 versus 0.64 genome-wide, **Supplementary Figure 13**). However, we found that only a small fraction (7.97% on average) of uncolocalized autoimmune GWAS loci showed evidence of a regulatory effect using this approach. This would still leave about half of all autoimmune GWAS loci uncolocalized.

To further test the possibility that the regulatory effects of many GWAS loci is outside current range of detection owing to small sample sizes, we asked how many uncolocalized GWAS loci could be colocalized using eQTL summary statistics from eQTLGen, which were obtained from a meta-analysis of 31,684 whole blood samples [41]. As quality control, we first compared the eQTLGen colocalizations with that of DGN (which sampled the same tissue type as eQTLGEN, whole blood, 15,269 common genes) and that of BLUEPRINT (15,373 common genes). Of the 242 autoimmune GWAS loci that colocalized with DGN eQTLs, 196 were found to replicate using the eQTLGen dataset (168 of 232 (72.4%) for BLUEPRINT). The higher replication rates for DGN was to be expected given that DGN and eQTLGen sampled the same tissue-type, whole blood, while BLUEPRINT assayed sorted immune cell-types.

Using eQTLGen eQTLs, we identified an additional 130 GWAS loci that colocalize in eQTLGen but not in BLUEPRINT, on average accounting for 16.8% (range: 6.6% - 35.8%) of uncolocalized loci from our BLUEPRINT analyses (**Supplementary Figure 14**). These findings suggest that although the gain in colocalization by increasing sample size could be large for some GWAS (e.g. 35.8% for multiple sclerosis), the average increase in colocalization rate is small. As expected, colocalized eGenes specific to eQTLGen tend to not be eGenes in DICE immune cell-types, or were eGenes with cell-type-specificity (**Figure 5e**). Additionally, the eQTLs of colocalized eGenes specific to eQTLGen have smaller effect sizes on average than that of colocalized eGenes specific to DGN, which in turn have smaller effects on average than colocalized eGenes that were identified to be shared in the DICE dataset (**Figure 5f**).

Despite the improvement in detection power afforded by the large eQTLGEN sample size, the average GWAS colocalization rates for eQTLs only increased slightly, from 22.9% using BLUEPRINT compared with 29.4% using eQTLGEN. Indeed, even when colocalized loci ascertained in DGN and eQTLGen are combined together, only an average of 35.8% GWAS loci colocalized with an eQTL. We thus conclude that increasing sample size is unlikely to drastically improve rates of colocalization, at least for cell-types that are well-represented in whole blood.

### Condition-specific profiles of H3K27ac in RA patients highlights context-dependent effects in RA pathogenesis

Finally, we hypothesized that the effects of some uncolocalized GWAS loci may be more readily interpretable in the context of the corresponding disease. While stimulation of immune cells *in vitro* may capture some important regulatory features reflecting disease state, we reasoned that studying immune cells sampled directly from autoimmune disease patients may better help understand the effects of uncolocalized GWAS loci. To this end, we focused specifically on rheumatoid arthritis (RA), an autoimmune disease that primarily affects synovium joints and is often associated with immune cell infiltration that lead to the build up of synovial fluid (SF) that can be collected from a joint aspiration [43].

To obtain regulatory profiles of cells in the context of RA, we first collected peripheral blood mononuclear cells (PBMC) from 6 RA patients and 4 healthy controls, as well as synovial fluid from the same RA patients. We then sorted B cells, CD4^+^ and CD8^+^ T cells, regulatory T cells and monocytes using flow cytometry (Methods), and profiled regions marked with H3K27ac using CUT&Tag (**Figure 6a**). Using these data, we identified regulatory regions and quantified their activity in 5 immune cell-types and 3 different immune contexts corresponding to the peripheral immune context in a healthy state, the peripheral immune context in the disease state, and the immune context at the active site of inflammation. More specifically, we mapped CUT&Tag 150bp paired-end reads onto the genome using Bowtie 2 [44] and identified peaks using MACS2 [45] for each sample separately. We then merged the peaks for all samples, by joining peaks that overlap, to obtain a single consensus peak set that was used for quality control and downstream analyses (**Supplementary File 6**).

**Figure 6.**
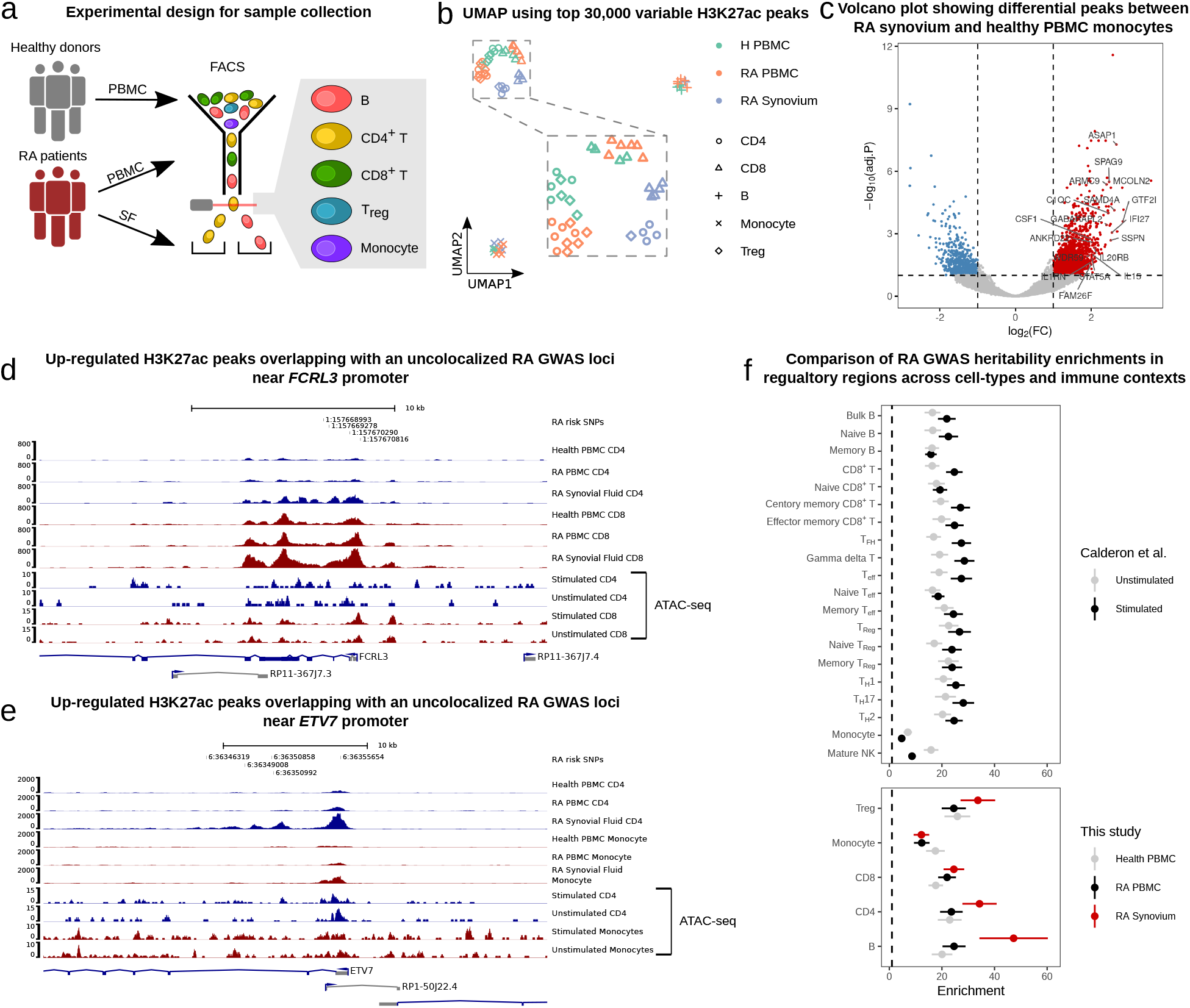
H3K27ac profiling in RA samples reveals disease-specific effects. **(a)** Schematic representation of our sample collection design. **(b)** UMAP of healthy and RA samples collected from PBMC and synovial fluid. The samples clustered by cell-type and by immune context. **(c)** Volcano plot showing differentially acetylated (H3K27ac) peaks between RA SF and healthy PBMC monocytes. **(d) and (e)** Examples of unexplained GWAS loci that overlap with regions that with higher H3K27ac activity in RA synovial fluid immune cells. RA risk SNPs were fine-mapped using SuSiE [42] and are shown along with their GWAS – log_10_ p-values. ATAC-seq peaks from Calderon et.al [39] were plotted for comparison. **(d)** H3K27ac activity at the *FCRL3* promoter is increased in RA SF CD4^+^ T cells (log_2_CPM: 4.02) compared to RA PBMC CD4^+^ T cells (log_2_-fold-change: 1.55, log_2_CPM: 2.46, FDR: 0.016) and healthy PBMC CD4^+^ T cells (log_2_-fold-change: 1.72, log_2_CPM: 2.30, FDR: 0.0077). For CD8^+^ T cells, the log_2_-fold-change is 1.32 compared to healthy PBMC. (e) H3K27ac activity at the *ETV7* promoter is increased in RA SF CD4^+^ T cells (log_2_CPM: 6.48) compared to RA PBMC CD4^+^ T cells (log_2_-fold-change: 1.89, log_2_CPM: 4.60, FDR: 0.0016) and healthy PBMC CD4 T cells (log_2_-fold-change: 2.47, log_2_CPM: 4.01, FDR: 5.70× 10^-5^). For monocytes, the log_2_-fold-change is 2.02 compared to healthy PBMC. **(f)** Forest plot of hetibability enrichment in ATAC-seq peaks (top) and H3K27ac CUT&Tag peaks from various cell-types (bottom) computed using stratified LDscore regression. RA heritability enrichments in H3K27ac peaks detected in T cells and B cells from RA synovial fluids are greater than that of ATAC-seq peaks of the same cell-types subject to *in vitro* stimulation. Error bars represent ± 1 standard error.

As expected, UMAP visualization of the log_2_-transformed read count-per-million (log_2_CPM) at the top 30,000 most variable peaks in the consensus set showed separation of the major cell groups (**Figure 6b**). In particular, B cells and monocytes formed distinct clusters, while CD4^+^, CD8^+^ and regulatory T cells clustered together. Notably, cells from the same biopsy site also formed subclusters such that immune cells from healthy and disease PBMC clustered more closely together, while immune cells from synovial fluid clustered separately. Importantly, samples did not cluster according to batch or other technical factors (**Supplementary Figure 15**), indicating that the observed clusters reflect biological differences between cell-types and biopsy sites.

We next compared H3K27ac activity between immune cells from the different immune contexts (Methods). The general trend we observed was that H3K27ac profiles in T cells were more different between RA synovial fluid and RA PBMC than between RA PBMC and healthy PBMC. Indeed, we found 5,355 and 6,339 differentially acetylated peaks between RA SF and RA PBMC cells for CD4^+^ T cells and CD8^+^ T cells, respectively, compared to the 1,045 and 1,070 differentially acetylated peaks between RA PBMC and healthy PBMC. By contrast, the H3K27ac profile of monocytes from RA PBMC is more similar to that of RA SF monocytes than that of healthy PBMC monocytes. This finding suggests that monocytes in the peripheral blood of RA patients show similar pathogenesis signatures to synovial fluid monocytes (e.g. at the *IL1B* locus, **Supplementary Figure 16**), and corroborates observations that were made previously using single-cell RNA-seq data [46].

We next studied the 5,958 peaks that showed higher activity in immune cells from RA SF compared to immune cells from healthy PBMC. We found that many of these peaks are located near important genes that are involved in inflammation pathways and disease pathogenesis, such as *CSF1,* which modulates the differentiation of monocytes to macrophages [47], and *IL1RN* (also known as *IL1RA),* which encodes the interleukin-1 receptor antagonist protein that has been associated with autoimmune diseases including RA [48]. Interestingly, *IL1RN* expression was also found to be upregulated in monocytes treated with synovial fluid from arthritic joints [49]. Overall, we found that genes near peaks with higher activity in RA SF monocytes were enriched in functional annotations such as immune response (P-value: 2.96 × 10^-14^, hypergeometric-test), immune effector process (1.76 × 10^-18^), and several pathways including interferon, TNF, NF-*κ*B, and TLR signaling pathways (1.64 × 10^-03^, 5.10 × 10^-05^, 3.49 × 10^-03^, 8.46 × 10^-03^, respectively) (Methods). Thus, the H3K27ac profiles of RA SF immune cells revealed elements that appear context-specific and likely relevant to RA pathogenesis.

We then asked whether differentially active peaks were enriched in unexplained GWAS loci. To answer this question, we overlapped differentially accessible peaks in all immune cells from RA patients with RA GWAS after fine-mapping using SuSiE [42] (Methods). Strikingly, we found that of the 42 uncolocalized RA GWAS loci, fine-mapped SNPs at 12 loci overlapped with a region with higher activity in RA immune cells (6 loci for healthy PBMC, bootstrap p-value 0.026, Methods). For example, we found that a lead GWAS SNP lies within a differentially active peak at the promoter region of *FCRL3* in CD4^+^ and CD8^+^ T cells (FDR: 7.7 × 10^-3^ for CD4^+^ and 2.6 × 10^-2^ for CD8^+^ T cells, **Figure 6d**). In another example, the RA lead SNP overlaps with an H3K27ac peak located near the promoter of *ETV7* which showed higher activity in both RA SF CD4^+^ T cells compared to the respective cell-types from RA PBMC (FDR: 1.6 × 10^-3^) and healthy PBMC (FDR: 5.7 × 10^-5^). The activity of this regulatory region was also higher in RA SF monocytes compared to healthy PBMC monocytes (FDR: 4.5 × 10^-3^, **Figure 6e**).

To further assess the relevance of each immune context on the study of disease etiology, we quantified the enrichment of RA heritability in H3K27ac peaks identified in the immune cell-types from the different immune contexts. To establish a baseline for comparison, we used stratified LDscore regression to estimate the RA heritability enrichment in published accessible chromatin regions identified using ATAC-seq data from unstimulated and stimulated immune cell-types [39] (Methods). Our estimates recapitulated the findings from the original study [39] in which CD8^+^ T cells and delta gamma T cells showed the largest increase in heritability enrichment subsequent to stimulation (~30-fold vs ~20-fold enrichments for stimulated versus unstimulated). We then applied the same LDscore regression analysis to our H3K27ac peaks. We found that while the estimated RA heritability was similarly enriched in ATAC-seq peaks from unstimulated immune cells and in H3K27ac peaks from RA PBMC and Healthy PMBC (~20-fold), the heritability enrichment was greater for B cells, CD4^+^ T cells, and Tregs in peaks from RA SF than in peaks from *in vitro* stimulation of the same cell-types (**Figure 6f**).

Our analyses therefore show that there are significant differences in the regulatory landscapes of immune cell-types across disease states and immune contexts. In particular, we found that the regulatory landscape of cells extracted from the active site of RA inflammation showed striking differences when compared to that of circulating immune cells in the periphery of both RA patients and healthy individuals. Importantly, we find that the regulatory regions identified in immune cells from RA synovial fluid overlap with many uncolocalized GWAS loci and are the most highly enriched in RA SNP heritability. Altogether, these observations indicate the importance of studying cell-types in the correct disease context in order to elucidate the genetic etiology of a disease.

## Discussion

The goal of this study was to establish a detailed accounting of the effects of genetic variants on gene regulation in immune cells and their overlap with genetic effects on human traits and disease. Recent studies suggested that fewer than half of GWAS loci colocalize with an eQTL [5, 13]. This finding implies that much is left to be understood about the mechanisms by which genetic variants impact human traits.

There are several possible explanations for the small fraction of GWAS loci that colocalize with an expression QTLs. Our work evaluated the possibilities that (i) there exist genetic effects on gene regulation other than steady state gene expression levels, (ii) genetic effects are often restricted to cell-types and cell-state that are causal for the trait, and (iii) genetic effects are often too small to be detected, even in the causal cell-types or cell-states. These possibilities are not mutually exclusive, but the implications are different for how we should design future human genomics research. For example, if trait-associated variants often impact mRNA splicing but not steady state mRNA expression, then a more widespread focus on mapping the effects of genetic variants on mRNA splicing is needed. If most disease-associated genetic effects are very specific to cell-types and cellstates that are relevant to the trait, then studying eQTLs identified in bulk, unsorted, tissues will have limited success in elucidating the mechanisms underlying most GWAS loci.

Using our harmonized regulatory QTL data, we found that eQTLs and sQTLs together colocalized with up to 45% of trait-associated loci for the 72 GWAS we analyzed. On average, 40.4% of significant loci from the 50 immune-related GWAS colocalized with a regulatory QTLs, a larger proportion compared to an average of 26.4% for the 21 non-immune GWAS we analyzed (excluding Alzheimer’s disease, 55.2% colocalized). One of the caveats in our colocalization analysis is the use of the method COLOC. Although COLOC is a very popular method for colocalization analyses, it uses priors that, when altered, can impact substantially the computed posterior probabilities that the causal eQTL and GWAS variants are the same variants. Reassuringly, when we used another colocalization method, HyPrColoc [50], that does not rely on user-defined priors, we were able to replicate nearly all colocalized genes identified using COLOC, indicating that our colocalization analyses are robust and replicable (Supplementary Note 4).

Our data also allowed us to ask whether regulatory QTLs are likely to be active in many immune cell-types, or only in few or a single cell-type. We found that at least 40% eQTLs (81% for sQTLs) are shared across all 15 immune cell-types we analyzed from the DICE dataset. For closely-related cell-types, we found that the fraction of shared eQTLs was as high as 96% (99% for sQTLs). Intriguingly, activated and naïve T cells share nearly 70% of detected eGenes. Thus, QTL effects appear similar across many cell-types and cell states. One important implication of this finding is that eQTLs that colocalize with GWAS SNPs in one cell-type are also likely to be active in other cell-types. Thus, eQTLs that colocalize at a GWAS locus in one cell-type should, in general, colocalize in the other cell-types. Indeed, after accounting for variability in the posterior probabilities of colocalization reported by COLOC owing to the inherent noise in QTL mapping, we found that the majority of GWAS loci colocalizes with the same QTL in multiple cell-types. Altogether, these data questions the notion that the vast majority GWAS SNPs affect gene regulation in a very cell-type-specific manner as highlighted in several studies [7, 11]. Thus, the use of regulatory QTLs from proxy cell-types or tissues, e.g. from the GTEx consortium, to identify causal genes may be well justified for a large fraction of genes.

A noteworthy finding from our colocalization analysis is that genetic variants that impact mRNA splicing often colocalize with a GWAS signal. Indeed, if we considered eQTLs only, our rates of colocalization would be very similar to that of previous studies (26.2% vs 21%) [13]. Instead, when sQTLs were tested for colocalization, we found that more GWAS loci colocalized with sQTLs than with eQTLs. It is worth noting however, that a substantial number of GWAS loci colocalized with both an eQTL and an sQTL. This may be due to horizontal pleiotropy, whereby a genetic locus can influence the expression level of a gene, as well as the splicing of an intron in the same or a different gene. Another possible explanation for this observation is that eQTL effects are often mediated by sQTLs or vice-versa. A colocalization analysis for sQTLs conditioned on the eQTLs would be necessary to tease apart these possibilities but is outside the scope of our work.

Despite a substantial increase in colocalization rates in our study, we find that for most traits, over half of all GWAS loci do not colocalize with a regulatory QTL. Interestingly, we found several differences between genes at colocalized GWAS loci and those at uncolocalized loci. Genes at GWAS loci without colocalized regulatory QTLs tend to be more lowly expressed, have higher enhancer density, and are less tolerant to loss-of-function mutations. These findings suggest that genes at uncolocalized GWAS loci may be subject to stronger constraints both at the levels of gene regulation and sequence conservation. Thus, a plausible explanation is that genetic effects at these loci are on average smaller and more cell-type or context-specific compared to genes at GWAS loci with colocalization. This hypothesis is consistent with the idea that much larger sample sizes may be required to find the causal QTL effects that explain the associations at GWAS loci without colocalization. That said, our colocalization analyses on QTL datasets with very large sample sizes (DGN: N = 900, eQTLGen: N = 31,684) revealed that the rates of colocalization only increased slightly despite the large increase in our power to detect low-effect QTLs. We speculate that an important reason for the modest increase in colocalization is because both DGN and eQTLGen QTL data are from whole blood samples, which are less likely to capture genetic effects that are cell-type or context-specific.

One intriguing finding from our analyses is that eQTLs identified specifically in *in vitro* stimulated immune cells from DICE colocalized with only a small number of GWAS loci that did not colocalize with QTLs from unstimulated cells. This observation might seem surprising because a recent paper showed that autoimmune disease SNP heritability is more highly enriched in accessible chromatin from *in vitro* stimulated immune cells compared to naïve immune cells [39]. However, we should note that our data does indeed suggest that SNPs at colocalized and uncolocalized GWAS loci are more highly enriched in open chromatin from stimulated cells compared to unstimulated cells. The differences in the enrichment, however, is negligible, suggesting that stimulation-specific effects can not explain why a large fraction of GWAS loci do not colocalize with the regulatory QTLs identified in our study.

One possible explanation for the modest increase in the colocalization rates, when using eQTLs identified in stimulated immune cells, is that the immune cells stimulated *in vitro* only partly recapitulate gene regulation in the *in vivo* disease context. Thus, although many regulatory elements are primed to be activated subsequent to *in vitro* stimuli – thereby capturing some of the important regulatory regions relevant to disease – they may require additional factors to fully capture the effects of genetic variants on gene expression levels in the disease context. In support of this, [51] found that stimulating immune cells *in vitro* was able to recapitulate gene expression signatures of immune cells from rheumatoid arthritis (RA) patients when 6 different cytokines were used together, but not when the cytokines were used on their own.

To better understand the role of context on our ability to interpret GWAS signals, we collected H3K27ac measurements in healthy and RA patients using CUT&Tag to use as proxy for enhancer and promoter activity. Although the sample sizes are too small for a QTL analysis, we were able to use these data to ask whether gene regulatory data in the disease context could aid us identify putative mechanisms that underlie RA GWAS hits, in particular for loci with no QTL colocalization. We found that SNPs at 12 out of 42 uncolocalized GWAS loci overlap with regions with increased H3K27ac levels in immune cells from RA synovial fluid. Remarkably, we also found that regions marked by H3K27ac in immune cells from RA synovial fluid were more highly enriched in RA heritability than compared to healthy or RA immune cells collected from peripheral blood. It would be interesting to perform a direct comparison between regulatory regions from *vitro* stimulated immune cells and that from RA synovial fluid immune cells. However, our initial analyses suggest that the RA GWAS heritability enrichments in regulatory regions identified in RA synovial fluid immune cells are higher than in that of *in vitro* stimulated immune cells. We should note here that caution must be used when interpreting these results as the data type collected in these two studies differ (ATAC-seq versus CUT&Tag). Nevertheless, these preliminary analyses indicate that studying the regulatory effects of genetic variants in the disease context may be critical for discovering the mechanisms behind a large number of GWAS loci.

## Methods

### Data processing

To harmonize the set of genetic variants across all four datasets, we imputed the genotypes of all individuals in the four studies using the 1000G Phase 3 v5 as a common reference panel (Michigan Imputation Server [52]). Following imputation, only non-duplicated genetic variants with INFO score larger than 0.9 were retained. We filtered variants with Hardy-Weinberg Equilibrium (HWE) p-values below 10^-5^, with missing genotype rate higher than 5%, and with minor allele frequency below 5% using PLINK v1.9 [53]. We used the remaining set of variants in all subsequent analyses unless otherwise noted. To exclude outlier individuals, we calculated genotype principal components (PCs) using smartpca [54]. Five outliers in the DICE dataset were identified and removed from downstream analyses.

To quantify gene expression levels, we used `Kallisto` [55] and summed the transcript per million (TPM) estimates of all GENCODE 19 [56] isoforms to obtain a gene-level TPM. The gene-level TPM were then scaled and quantile-quantile normalized as described before [57]. Gene expression principal components were calculated using the `prcomp` function in `R.` To quantify RNA splicing, RNA-seq reads were aligned to the hg19 reference gnome using `STAR 2.6.0` [58] with the GENCODE 19 annotation. To avoid reads mapping with allelic bias, we used `WASP` [59] as implemented in `STAR 2.6.0` by providing the corresponding genotype data. This is an important step as we found a substantial increase in the number of false positive splicing QTL due to allelic bias in read mapping [20]. Exon-exon junctions were extracted using `RegTools` [60], and clustered and quantified using `LeafCutter` [20]. As expected, we observed that the number of exon-exon junctions identified in each sample is positively correlated with the sequencing depth in the DICE consortium (**Supplementary Figure 1**). To harmonize quantification for splicing junction usage across cell-types and datasets in all 18 immune cell-types, clusters were merged and the merged union was used to re-calculate intron usage in all samples.

### MashR analysis in the DICE dataset

To quantify the sharing of eQTLs and sQTLs in the DICE dataset, we followed the workflow provided by the authors of MashR (https://github.com/stephenslab/gtexresults) that was previously described in [18]. Briefly, standard errors of QTL effect sizes were calculated from `FastQTL` nominal output, which were used together with effect sizes as the input for *mash.* To obtain a confident set of QTLs for each feature (gene or intron), the SNP with the smallest P-value across all tested SNPs and all cell-types were extracted for each feature. This resulted in a feature-by-sample matrix of effect sizes and their standard errors without missing values. For eQTLs, we included all protein coding genes for training. For sQTLs, we included all introns for training. We then built a *mash* model using the exchange effects (EE) mode to estimate the priors. This model was then applied to all QTLs to calculate the posterior mean effect sizes (*mash* effect sizes). Significant QTLs after *mash* analysis were feature-SNP pairs with local false sign rate (LFSR) below 0.05, as suggested by [18]. The level of QTLs sharing was quantified as both overall sharing and pairwise sharing. Overall, sharing was determined to be the number of cell-types in which a given feature has a regulatory QTL (LFSR < 0.05). Pairwise sharing was quantified both by magnitude and by sign. Sharing-by-magnitude between two cell-types correspond to the proportion of QTLs that is significant in one of the cell-types and posterior mean effect sizes differ by no more than two-fold. Sharing-by-sign between two cell-types correspond to the proportion of QTLs that was significant in one of the cell-types and had the same sign. The 15 cell-types in DICE were grouped into 6 cell-groups based on the eQTL sharing-by-magnitude (see **Figure 2b**).

### Characterization of regulatory QTLs

To calculate the distance between eQTLs and their target genes, we defined the promoter of each gene as the region 2,000bp upstream and 500bp downstream of TSS. We tested the enrichment of eQTLs in regulatory elements from Ensembl Regulatory Build and consensus ATAC-seq peak set from Calderon *et al.* [39]. We categorized all ATAC-seq peaks to be either an enhancer or a promoter based on whether they overlap with any promoter region (2,000bp upstream and 500bp downstream of TSS). The observed and expected number of QTLs overlapping with each feature was estimated using the `fenrich` command from `QTLtools` [61], and the odds ratios of enrichment were calculated by supplying those number to Fisher’s exact test in R. We validated eQTLs from DICE in other datasets using *π*_1_ statistics [62], stratifying eQTLs by their levels of sharing across six cell-groups estimated by *mash* (specific: in one cell-group; intermediate: 2–5 cell groups; shared: 6 cell-groups). The 95% confidence intervals of *π*_1_ was estimated using 1,000 bootstraps (i.e. resampling DICE eQTLs with replacement).

### Colocalization

#### COLOC

Colocalization analyses were performed between eQTLs/sQTLs and 72 publicly available GWAS summary statistics for 11 autoimmune diseases (14 studies), namely, rheumatoid arthritis (RA) [63], Crohn’s disease (CD) [23, 26], ulcerative colitis (UC) [23, 26], inflammatory bowel disease (IBD) [23, 26], allergy and eczema (AE) [64], asthma, hay fever and eczema (allergy for short) [65], apoptotic dermatitis (ApD) [66], asthma [67, 68], systemic lupus erythematosus (SLE) [69] and multiple sclerosis [70]. We also collected 36 GWAS for blood-related traits [71], 11 GWAS related to heart functions and circulation system [72], and several other traits including type 2 diabetes (T2D) [73], Alzheimer’s disease (AD) [74], Parkinson’s disease (PD) [75], estimated glomerular filtration rate (eGFR) [76], height [77], and breast cancer survival [78] and other cancers/neoplasms [72]. We considered the 14 autoimmune and the 36 blood-related GWAS as immune GWAS, and the rest 22 GWAS as non-immune GWAS.

To assess colocalization between GWAS loci and QTLs, we first identified the lead GWAS variants and their flanking region in which colocalization was to be tested. Specifically, all variants available in the GWAS summary statistics were sorted by p-values in increasing order. Starting from the variant with the smallest p-value (lead variant), variants within the 500Kb window on either side of the leading variant were removed. This resulted in a 1Mbp GWAS locus for colocalization analysis. The same procedure was then applied to the next most significant variant among the remaining variants, until no variant with p-value below 10^-7^ was left. The HLA region (Chr6: 25Mb-35Mb) was excluded from colocalization. Only GWAS with more than 10 identified loci were included in our analysis. For each GWAS locus identified above, colocalization was tested only if it harbored a regulatory QTL with beta-distribution permuted p-value below 0.01 (bpval < 0.01) as reported by `FastQTL` in the 1Mb window flanking that leading GWAS SNP. Default priors were used for `COLOC`. We set PP4 > 0.75 as the threshold for colocalization. The colocalization proportion was calculated as the proportion of colocalized loci among all identified loci in a GWAS.

Colocalization results were visualized using a function adapted from LocusCompare [79]. For a given locus, SNP with the largest posterior probability from `COLOC` was defined as the colocalized SNP. *r*^2^ relative to the colocalized SNP were calculated from the genotypes in the QTL study.

To visualize the sQTL in the form of a Sashimi plot [80], we first grouped individuals by their genotypes, and then extracted RNA-seq reads that mapped to the cluster that contains the intron to be visualized. To make the coverage comparable between different genotypes, we scaled the read coverage by the number of indivuduals that carry each genotype using the `scaleFactor` argument in `bamCoverage` from Deeptools [81] when generating bigWig files. The coverage was then visualized using pyGenomeTracks [82].

Cis-eQTL data of eQTLGen [41] was directly obtained from the website (https://eqtlgen.org/cis-eqtls.html). We also downloaded allele frequencies from 26,609 eQTLGen samples (excluding Framingham Heart Study), which were used in our colocalization analysis.

#### HyPrColoc

The GWAS-gene pairs tested in `HyPrColoc` were selected in the same way as `COLOC`. We set PP > 0.25 as the threshold for colocalization as recommended by the authors [50].

#### Validation of immune-cell-specific colocalization for non-immune traits

We validated colocalization of 14 non-immune traits (11 heart-related, AD, PD and breast cancer survival) in DICE immune cells using the GTEx V7 eQTLs. We first chose several tissues in GTEx that are most relevant to each GWAS trait. For heart-related traits, we chose tissues in heart and circulation system (Artery - Aorta, Artery - Coronary, Artery - Tibial, Heart - Atrial Appendage, Heart - Left Ventricle). For AD and PD, we included the 13 brain tissues (Brain - Amygdala, Brain - Anterior cingulate cortex (BA24), Brain - Caudate (basal ganglia), Brain - Cerebellar Hemisphere, Brain - Cerebellum, Brain - Cortex, Brain - Frontal Cortex (BA9), Brain - Hippocampus, Brain - Hypothalamus, Brain - Nucleus accumbens (basal ganglia), Brain - Putamen (basal ganglia), Brain - Spinal cord (cervical c-1), Brain - Substantia nigra). For breast cancer survival, we used adipose tissues and breast tissue (Adipose - Subcutaneous, Adipose - Visceral (Omentum), Breast - Mammary Tissue). We then identified all the colocalized gene-SNP pairs for these 14 GWAS in DICE, and extracted their P-values from GTEx eQTLs in the relevant tissues, as well as from DICE eQTLs in all immune cell-types. Given that a large proportion of eQTLs are shared in DICE, we grouped the 15 immune cell-types into 6 groups, assigning the smallest P-value from all cell-types within a given group to that group for each gene. We used Bonferroni correction to adjust P-values for multiple testing. Finally, we calculated the proportion gene-SNP pair that has adjusted P-value below 0.05 in DICE but not GTEx tissues.

#### Characterizations of uncolocalized GWAS loci

We restricted this analysis to the loci from the 14 autoimmune GWAS that did not colocalize with a in BLUEPRINT QTL. All genes were classified into four categories: genes with an eQTL that colocalized at a GWAS locus, genes that are the closest to a GWAS locus, genes that are closest to a uncolocalized GWAS locus, and all remaining genes. We compared gene expression level in the three BLUEPRINT cell-types separately. The gene expression level values for the three cell-types were combined and plotted in **Figure 5a**. We also obtained Enhancer-domain score (EDS) [38] and “loss-of-function observed/expected upper bound fraction” (LOEUF) [36] for all available genes and compared the distribution of EDS and LOEUF across the four categories above.

To test the enrichment of uncolocalized loci in ATAC-seq peaks in stimulated immune cells, we constructed a contingency table by counting the number of colocalized and uncolocalized loci overlapping stimulated and unstimulated ATAC-seq peaks, respectively. We then tested the hypothesis that uncolocalized loci were more highly enriched in stimulated open chromatin regions compared to colocalized loci using Fisher’s exact test. We estimated 95% confidential interval of estimates by bootstrapping uncolocalized GWAS loci 1,000 times with replacement.

We reasoned that regulatory effects of many uncolocalized GWAS loci might be too small to be detected due to small sample sizes. To test this possibility, we ascertained eQTLs only at uncolocalized GWAS loci. Briefly, we extracted QTL tests at lead SNP of uncolocalized loci. GWAS locus-gene pairs that have already been tested in `COLOC` but did not colocalize were filtered. Since it is common for one lead SNP to be associated with many genes, we adjusted the P-values by number of tested genes at each loci using Bonferroni correction and picked the gene with the smallest P-value. We then calculated the proportion of genes with P-value below 0.05. This analysis was applied to each autoimmune GWAS in each cell-type in BLUEPRINT dataset.

### RA samples collection and analysis

#### Sample collection and CUT&Tag experiment

The collection of human samples was approved by the Institutional Review Board of the Xijing Hospital (Xi’an, China) and has obtained written informed consent from all participants. All of the clinical samples were obtained from Xijing Hospital. Peripheral blood and synovial fluid samples were collected from 6 RA patients at the Department of Clinical Immunology, Xijing Hospital. All of the RA patients fulfilled the 1987 revised American College of Rheumatology criteria and the ACR 2010 Rheumatoid Arthritis classification criteria [83], and their clinical characteristics are shown in **Supplementary Table**. In addition, peripheral blood samples were gathered from 4 healthy individuals. All blood and synovial fluid samples were subjected to gradient centrifugation using lymphocyte separation medium (MP Biomedicals, 0850494) to isolate mononuclear cells, which were cryopreserved for later experiments.

The cryopreserved mononuclear cells were thawed into RPMI/10%FBS, washed once in sterile phosphate-buffered saline (PBS; Beyotime, ST476), and stained with the following antibodies in PBS for 30 minitues: anti-CD3-APC/Cy7 (Biolegend, 300426), anti-CD4-PE/Cy7 (Biolegend, 357410), anti-CD8-Percp/Cy5.5 (Biolegend, 301032), anti-CD25-PE/CF594(BD Horizon,562525), anti-CD19-FITC (Biolegend,302206), and anti-CD14-APC (Biolegend, 301808). CD4^+^ T cells (CD3^+^, CD4^+^, CD8^-^), CD8^+^ T cells(CD3^+^, CD4^-^, CD8^+^), T_reg_ cells (CD3^+^, CD4^+^, CD8^-^, CD25^+^), B cells (CD3^-^, CD19^+^), and monocytes (CD3^-^, CD14^+^) were sorted by FACSAria III (BD Pharmingen, San Diego, USA) directly into wash buffer for CUT&Tag, with a maximum of 1 × 10^5^ cells for each cell type. We profiled H3K27ac (abcam ab4729) for each cell-type following standard CUT&Tag protocol (https://www.protocols.io/view/bench-top-cut-amp-tag-z6hf9b6) [17]. Samples were processed in different batches, and we ensured to include at least one healthy individual and one RA patient in each batch to minimize batch effects that align with biological differences that we are interested in.

#### CUT&Tag data analysis

The DNA libraries were subjected to 150bp paired-end (PE) sequencing. Sequencing reads were aligned to human reference genome hg19 using Bowtie 2 [44] with parameters `--local --very-sensitive-local --no-unal --no-mixed --no-discordant --phred33 --minins 10 --maxins 700 -p 16`. Aligned reads were filtered using Samtools with `-F 1804 -f 2 -q 30`. Samples with fewer than 2M reads were excluded from subsequent analyses. Filtered BAM files for samples that have the same disease status (healthy/RA), tissuetype (PBMC/SF) and cell-type were merged. Read coverage was calculated using `bamCoverage` in 10bp window normalized by RPKM [81]. H3K27ac peaks were called from the merged BAM files using `MACS2` with parameters `--format BAMPE --broad --broad-cutoff 0.1 --qvalue 0.1 --extsize 146` [45]. We reasoned that calling peaks from merged BAM files increases the signal-to-noise ratio. To generate a consensus peak set, we merged all the peaks using `bedtools merge`, resulting in 90,412 peaks. We then counted the number of fragments overlapping with the consensus peak set in each sample using `featureCounts` [84].

Differential peak analysis was performed using `limma` [85]. We calculated average log_2_CPM across samples with the same disease status, tissue-type, and cell-type. This average log_2_CPM was only used to filter our peaks with low fragments counts. Peaks with average log_2_CPM below 2 in all groups were excluded from differential analysis. Then, normalization factors were calculated from the remaining peaks using the TMM method, and counts in each sample converted to log_2_CPM. Since samples were processed in different batches, we used `ComBat` to adjust for batches while including disease status, tissue-type, and cell-type as our variable-of-interest. We constructed a contrast matrix comparing RA SF vs. RA PBMC, RA SF vs. Healthy PBMC, and RA PBMC vs. Healthy PBMC in each cell-type, and applied the `trend` method. Differential peaks were defined as log_2_-fold-change (log_2_(FC)) larger than 1 or smaller than −1, and FDR below 0.1.

We overlapped H3K27ac peaks up-regulated in RA samples with uncolocalized RA GWAS loci. We first fine-mapped RA GWAS summary statistics using `SuSiE` [42]. Fine-mapping was performed at each locus we used in our colocalization analysis. We supplied GWAS Z-scores, genotype correlation matrix from CEU and GBR from the 1000 Genome Project as the reference panel and the sample size of reference panel to the `susie_rss` function.

We estimated the enrichment of RA SNP heritability in our H3K27ac peaks using Stratified LD Score Regression (S-LDSC) [4]. We used MACS2 peaks from merged BAM files, which were extended by 500bp on both sides. To reproduce the heritability analysis from Calderon *et al.* [39], we used the MACS2 peaks shared by the authors.

## Data availability

All the Supplementary Files in the manuscript can be accessed through Box with: https://tinyurl.com/immuneqtl.

## Acknowledgement

We thank D Calderon, BE Mittleman, A Shah, F Morgante, and W Lin for comments on the manuscript. This work was supported by the US National Institutes of Health (R01GM130738 to YIL, R01CA242929 to Z Mu). This work was completed in part with resources provided by the University of Chicago Research Computing Center.

## Notes

https://tinyurl.com/immuneqtl

### Competing Interest Statement

The authors have declared no competing interest.

